## Supplementary figures and images for "Muscle-derived Myoglianin regulates *Drosophila* imaginal disc growth"

### Sup Fig 1 linked to Fig 1

Figure 1 Supplement 1

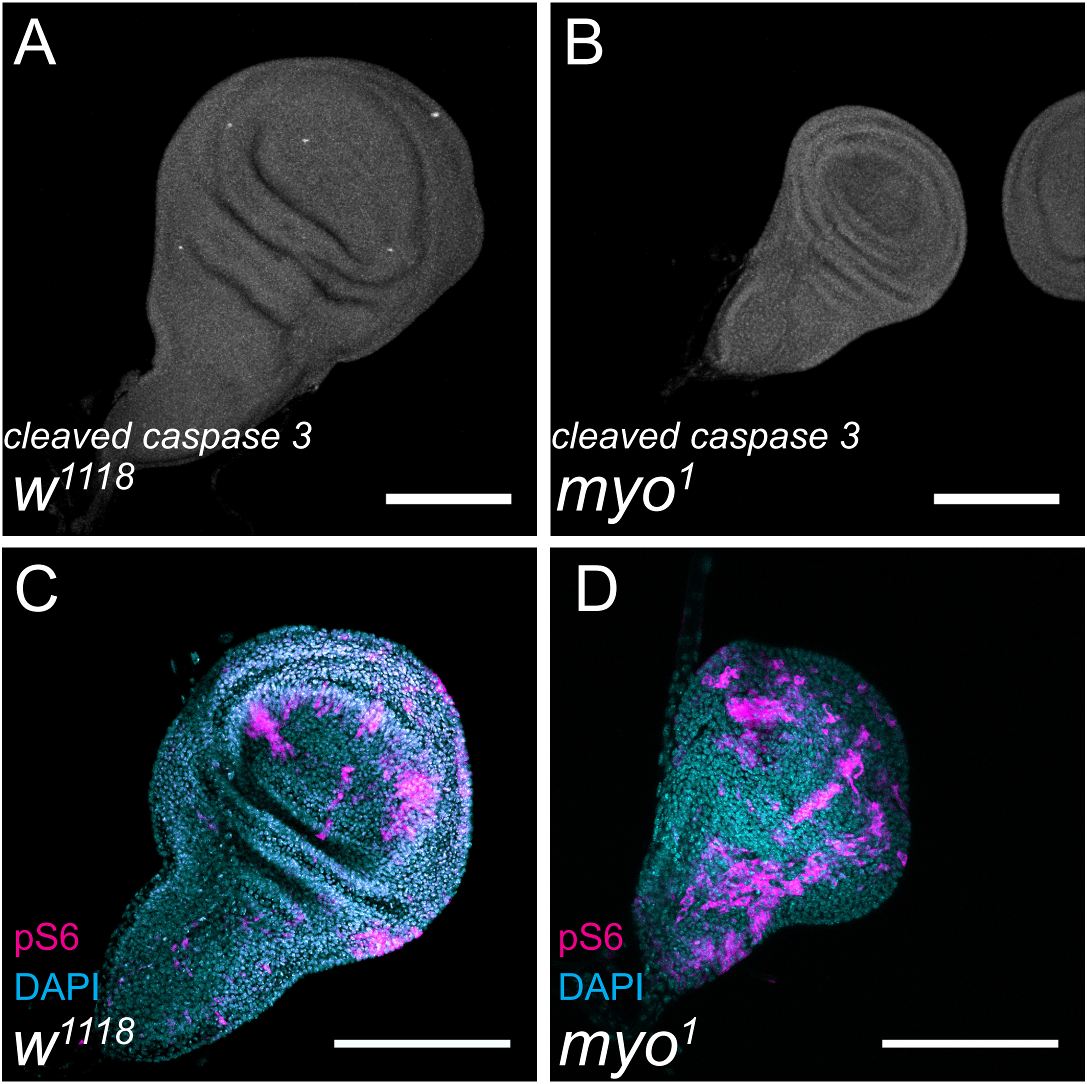

### Sup Fig 1 linked to Fig 2

#### Figure 2 Supplement 1

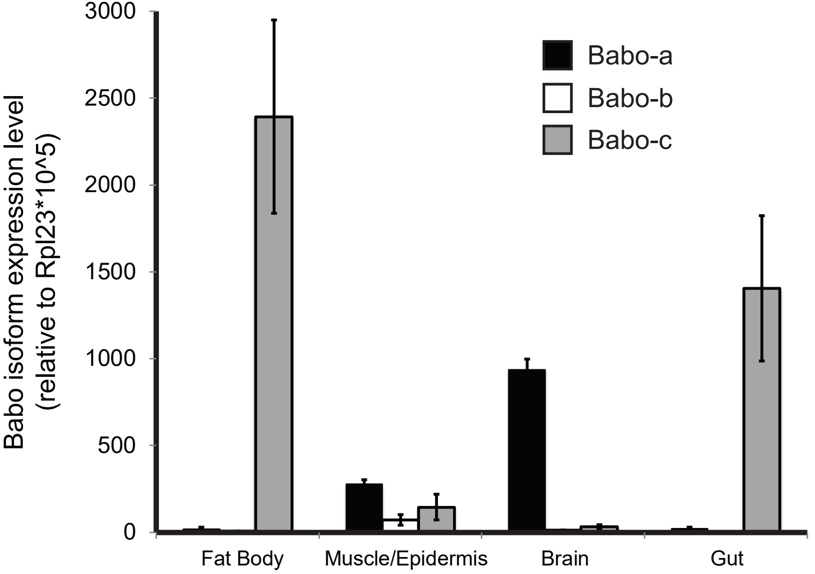

### Sup Fig 1 linked to Fig 3

Figure 3 Supplement 1

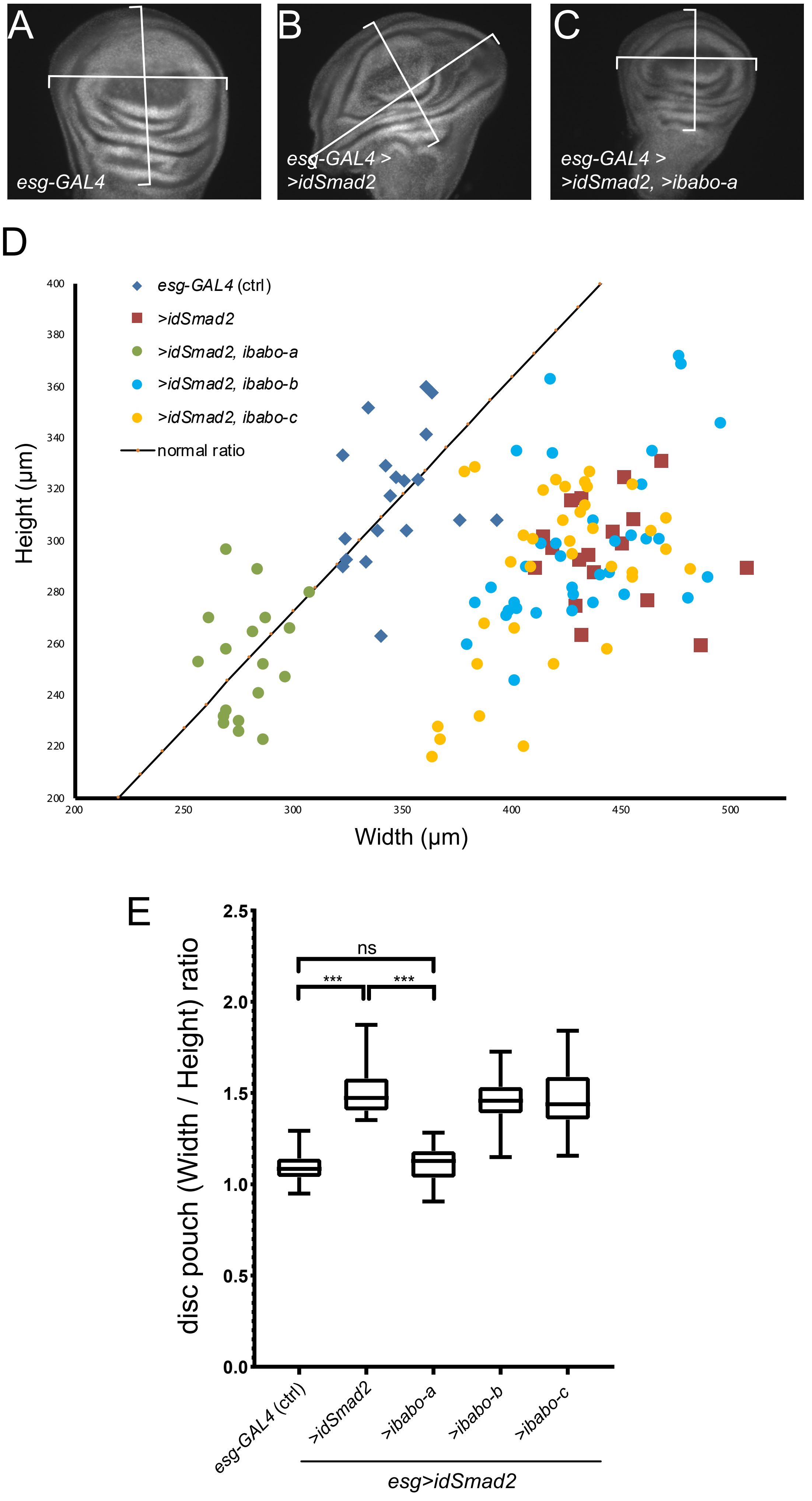

### Sup Fig 1 linked to Fig 4

Figure 4 Supplement 1

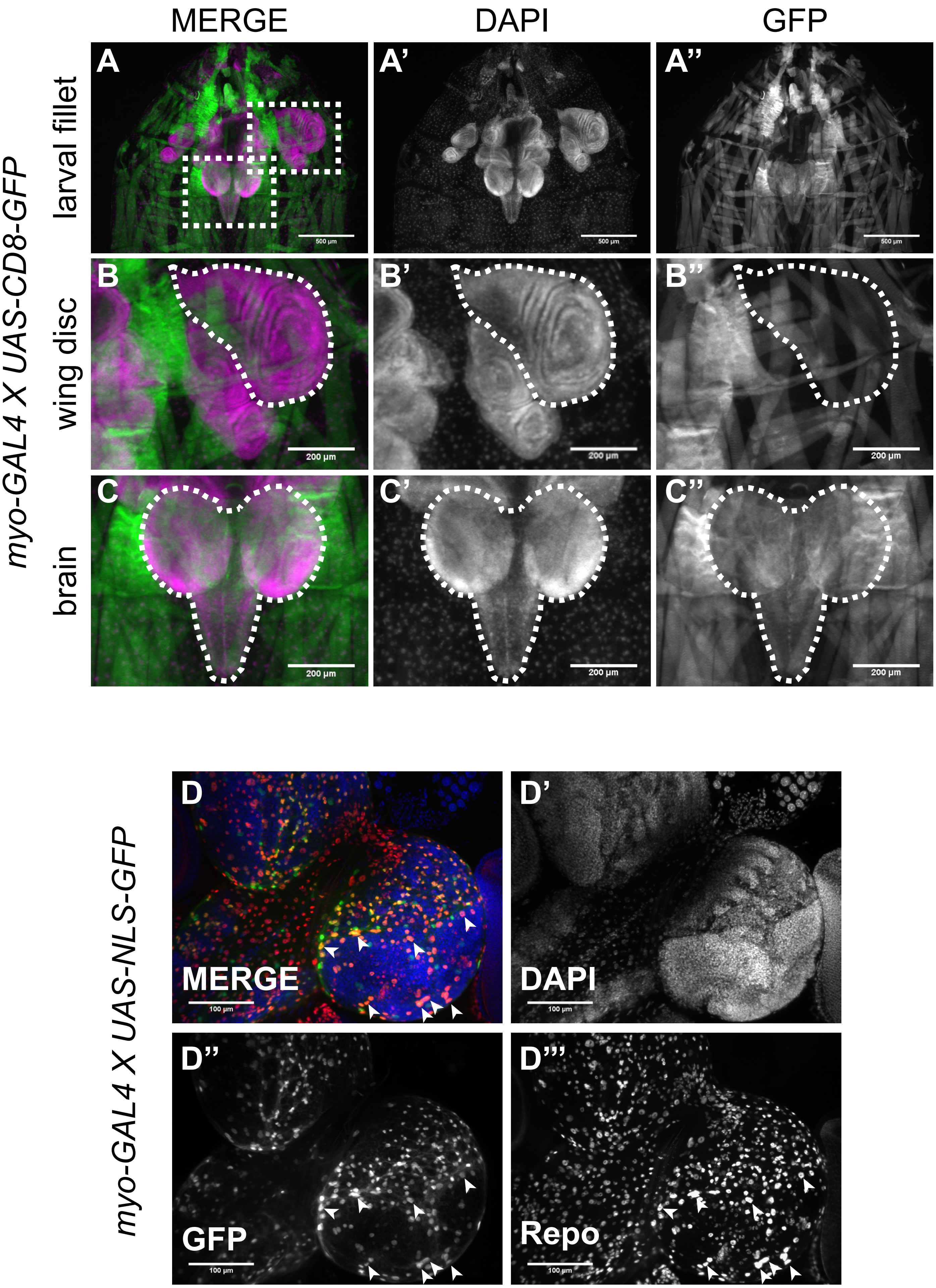

### Sup Fig 1 linked to Fig 5

Figure 5 Supplement 1

A

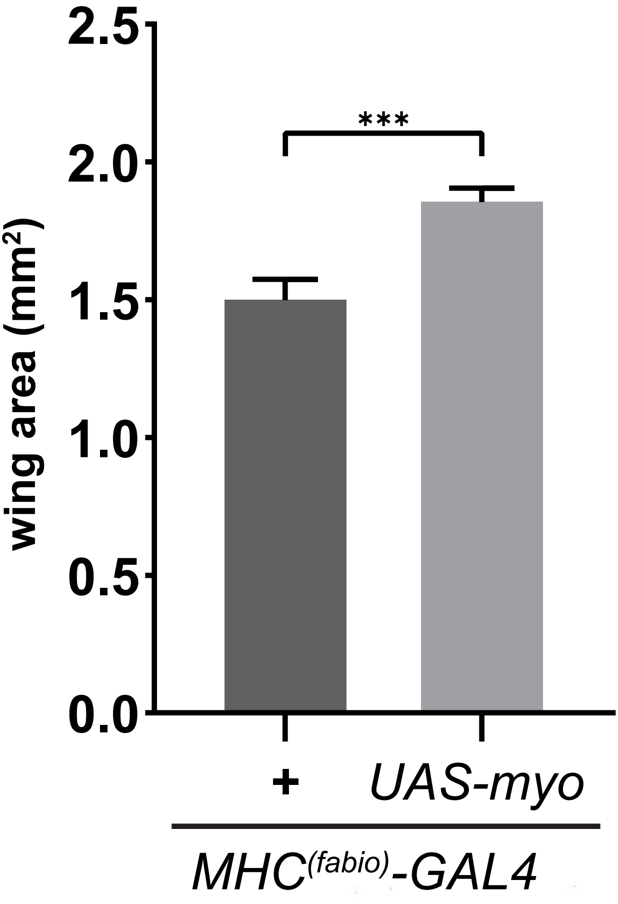


B

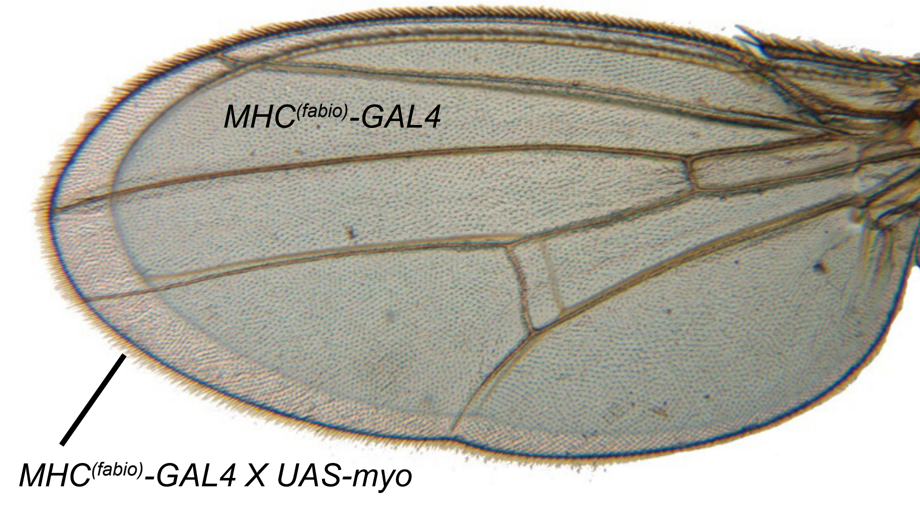

### Sup Fig 2 linked to Fig 1

Figure 1 Supplement 2

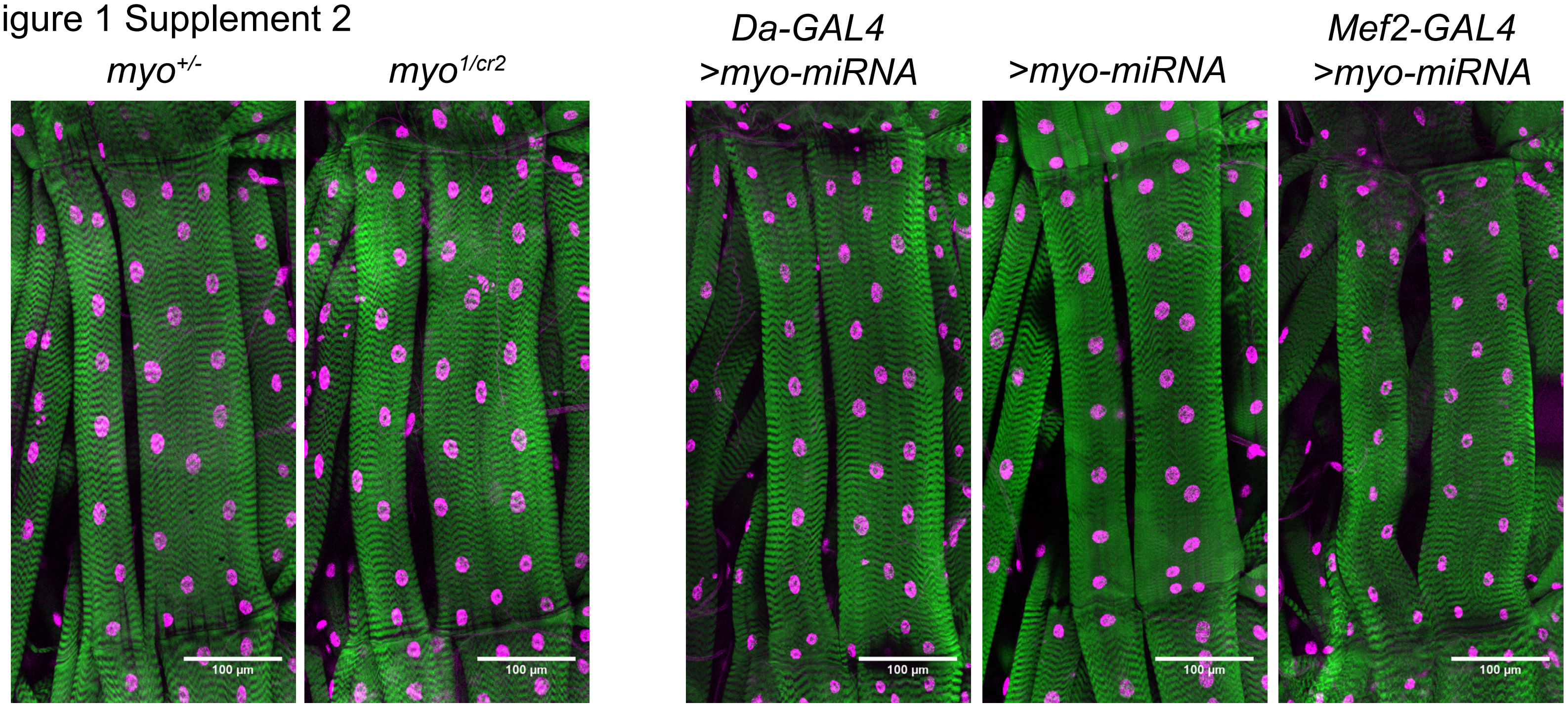


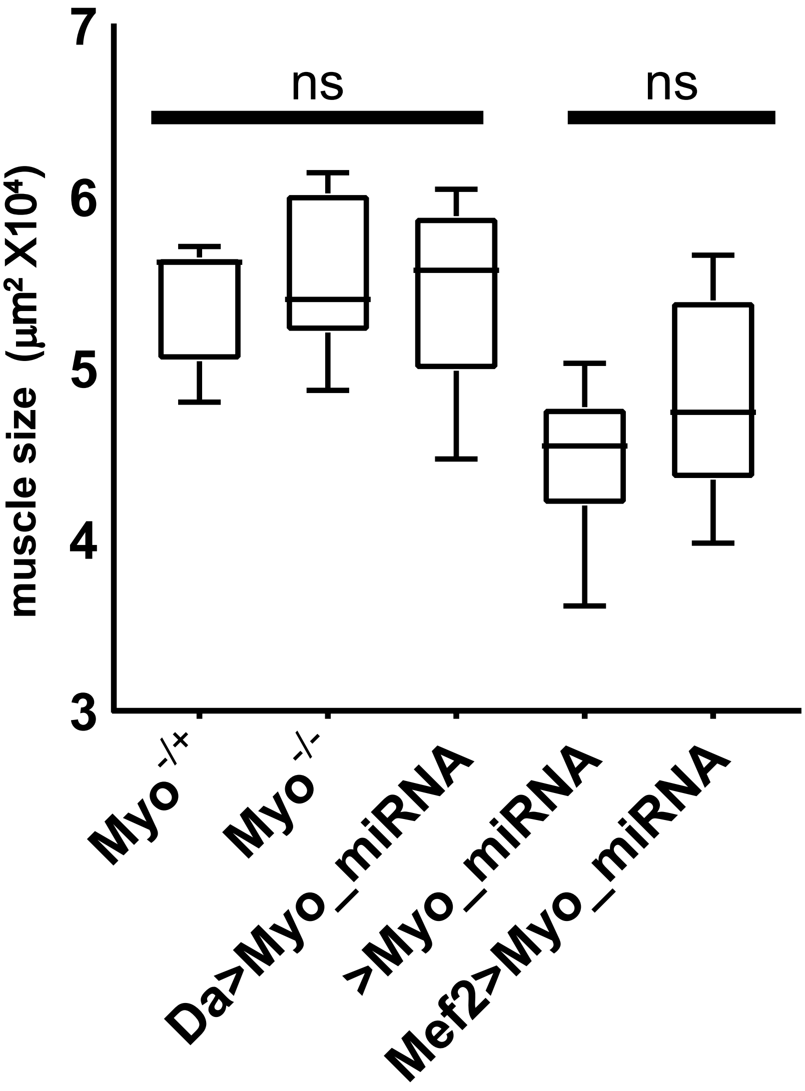
